# Fold first, ask later: structure-informed function annotation of *Pseudomonas* phage proteins

**DOI:** 10.1101/2025.07.17.665397

**Authors:** Hannelore Longin, George Bouras, Susanna R. Grigson, Robert A. Edwards, Hanne Hendrix, Rob Lavigne, Vera van Noort

## Abstract

Phages, the viruses of bacteria, harbor an incredibly diverse repertoire of proteins capable of manipulating their bacterial hosts, inspiring many medical and biotechnological applications. However, to date, only a limited subset of that repertoire can be exploited, due to the difficulties in functionally elucidating these proteins. In this study, we investigated several structure-informed approaches to annotate hypothetical proteins from *Pseudomonas* infecting phages. We curated a representative dataset of over 10,000 proteins derived from NCBI, for which we predicted protein structures with ColabFold and assessed structural similarity via FoldSeek against the PDB, AlphaFold, and Phold databases. We evaluated multiple annotation strategies, including sequence-based (Pharokka), and structure-based (FoldSeek, Phold) methods. Our results show that up to 43 % of truly unannotated proteins can be functionally annotated when combining structure-informed approaches with UniProt-derived annotations. We highlight the complementarity of different databases and the importance of annotation quality filtering. This work provides a valuable resource of predicted structures and annotations, and offers insights into optimizing structure-based annotation pipelines for viral proteins, paving the way for deeper exploration of phage biology and its applications.

## Introduction

Bacteriophages (hereafter, phages), the viruses of bacteria, are the most abundant creatures on earth^1^. They play a pivotal role in shaping microbial environments, and have been of critical importance to our understanding of molecular (micro)biology^2,3^. Moreover, phages and their proteins have inspired major biotechnological and medical applications^2,4,5^. Despite these clinical, biotechnological and fundamental interests in further deciphering phage biology, a vast amount of unknowns remain. Indeed, over 70% of open reading frames on phage genomes encode proteins of unknown function^6,7^. This “viral dark matter” stems from the significant genetic diversity harbored by phage proteins, hampering sequence similarity based annotation efforts^8^.

Fortunately, protein structure can often capture conserved functional relationships where sequence information falls short^9^. In the past, structural information was often unattainable due to the limited availability of protein structures. However, the release of AlphaFold2^10^ (and subsequent algorithms^11,12^) has virtually eliminated this bottleneck. Initially, viral proteins were excluded from large scale structure prediction efforts, such as the AlphaFold database^13^. Since then, viral protein structure databases have been emerging, including the phage protein containing BFVD^14^ and Viral AlphaFold Database^15^, and the phage specific PHROG structure resource^16^. While instrumental, the BFVD and the Viral AlphaFold Database have, in their efforts to maximally capture the viral protein space, grouped viral proteins far too loosely to be of direct use for function annotation.

To date, the only publicly available phage-oriented structure-informed protein annotation algorithm is Phold^17^. Phold uses the ProstT5 protein language model (pLM)^18^ to cast amino acids to FoldSeek-compatible 3Di tokens, bypassing the computationally expensive protein structure prediction process. These tokens are then used by FoldSeek^19^ to assess structural similarity to a curated set of phage protein structure databases^20–27^. To obtain a function annotation from the FoldSeek results, Phold uses a best annotated hit method.

Here, we assess the efficacy of structure-informed annotations, and the impact of algorithmic design choices when using FoldSeek for phage protein annotation. We focus on annotating hypothetical proteins of *Pseudomonas* infecting phages. This subset of phages was selected for three main reasons: (i) broad potential implications of results, given their biotechnological and medical relevance^28^, (ii) in-house expertise, aiding in the assessment of annotations, and (iii) computational feasibility. Briefly, we extracted all hypothetical proteins from NCBI protein^29^, and clustered them at 90% sequence identity and 80% coverage. Protein structures of cluster representatives were predicted with ColabFold^23^, and structural similarity to the PDB^30^ and AlphaFold database^13^ was assessed using FoldSeek^19^. Protein annotations were obtained from the FoldSeek output, following different post-processing strategies, as well as through Phold^17^ (and Pharokka^31^, as a structure-independent baseline). Following assessment of the different annotations, the data was reorganized in a phage-centric fashion, to provide easily accessible structural and functional information on the proteins.

## Results

### 71% of proteins from *Pseudomonas* infecting phages in NCBI protein are labeled ‘hypothetical/phage protein’

In February 2024, the Virus-Host database^32^ contained 887 phages that are (i) reported to infect *Pseudomonas* species and (ii) link to NCBI protein^29^. The majority of the dataset consists of dsDNA phages, mainly infecting *Pseudomonas aeruginosa* (98 % and 65 % respectively, Fig. 1A and B). At the time, 121,654 proteins could be retrieved from NCBI protein, 86,626 of which were labeled ‘hypothetical/phage protein’. At the level of individual phages, the unannotated proteins account for 67 ± 18 % of all proteins, which is weakly correlated to the total number of proteins (Pearson’s R^2^ = 0.06, Fig. 1C). Unannotated proteins were significantly shorter than their annotated counterparts, although length ranges overlap greatly (159 ± 166 vs. 343 ± 288 amino acids, p = 0.0, Fig. 1D).

**Fig. 1.**
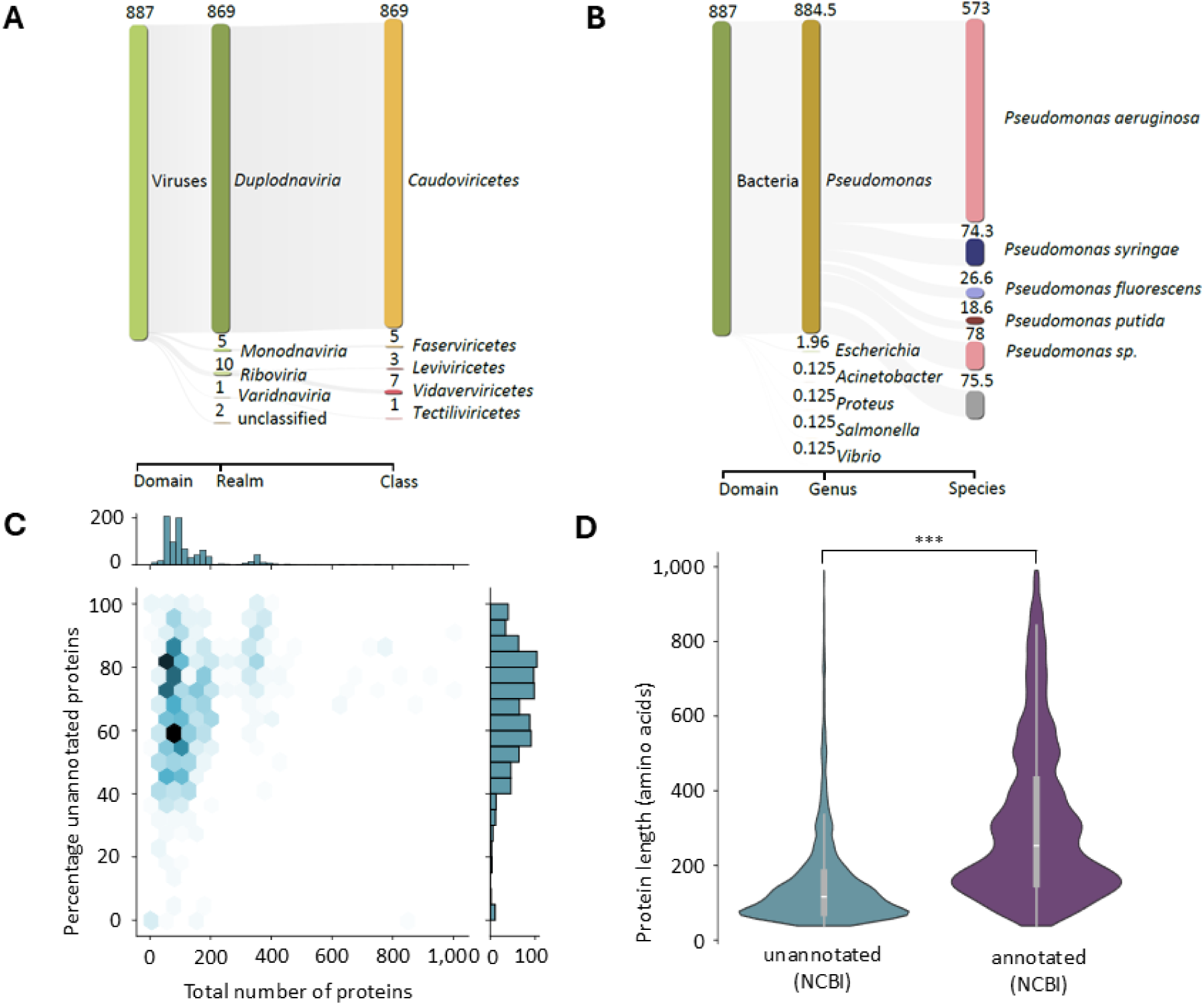
NCBI protein dataset statistics. **A. Phage taxonomy** of *Pseudomonas* infecting phages extracted from the Virus-Host database^32^. Shown at each level are the 5 most abundant taxa. **B. Host taxonomy** of *Pseudomonas* infecting phages extracted from the Virus-Host database. Phages with multiple recorded hosts were split proportionally. Shown at each level are the 6 most abundant taxa. **C. Joint distribution of the percentage of unannotated proteins and the total number of proteins** extracted from NCBI protein. Proteins were considered unannotated when the annotation contained the term ‘hypothetical protein’ and/or ‘phage protein’. **D. Distribution of protein length of unannotated and annotated proteins** extracted from NCBI protein. Violin plot excludes the 1% most extreme values on both ends. *** : p-value of 0.0 (one-sided Mann-Whitney U test).

Prior to structure prediction and annotation, the dataset of ‘hypothetical/phage’ proteins was further reduced: proteins were deduplicated, size filtered and clustered with MMseqs2^33^ (90% sequence identity, 80% bidirectional coverage). The final dataset consisted of 10,498 clusters, with an average cluster size of 4 proteins (average including the 6,978 singletons).

### Pharokka annotates 27 % of all ‘hypothetical/phage protein’ clusters using only sequence information

It should be noted that not all proteins in our dataset are necessarily truly unannotated. Indeed, due to the extraction strategy (most importantly: the use of redundant protein database NCBI protein), it is possible that proteins can be annotated in absence of structural information. This hypothesis was confirmed by Pharokka^31^, which annotated 27 % of cluster representatives (representing 30 % of all proteins). Most commonly, the Pharokka annotations pointed to structural proteins, as well as those related to nucleotide metabolism (Fig. 2A and 2B). Cluster representatives annotated by Pharokka belonged to slightly larger clusters than those who did not (hereafter referred to as ‘truly unannotated’, p = 3.4 x 10^-21^), and these proteins were significantly larger (p = 1.4 x 10^-212^, Fig. 2C and 2D).

**Fig. 2.**
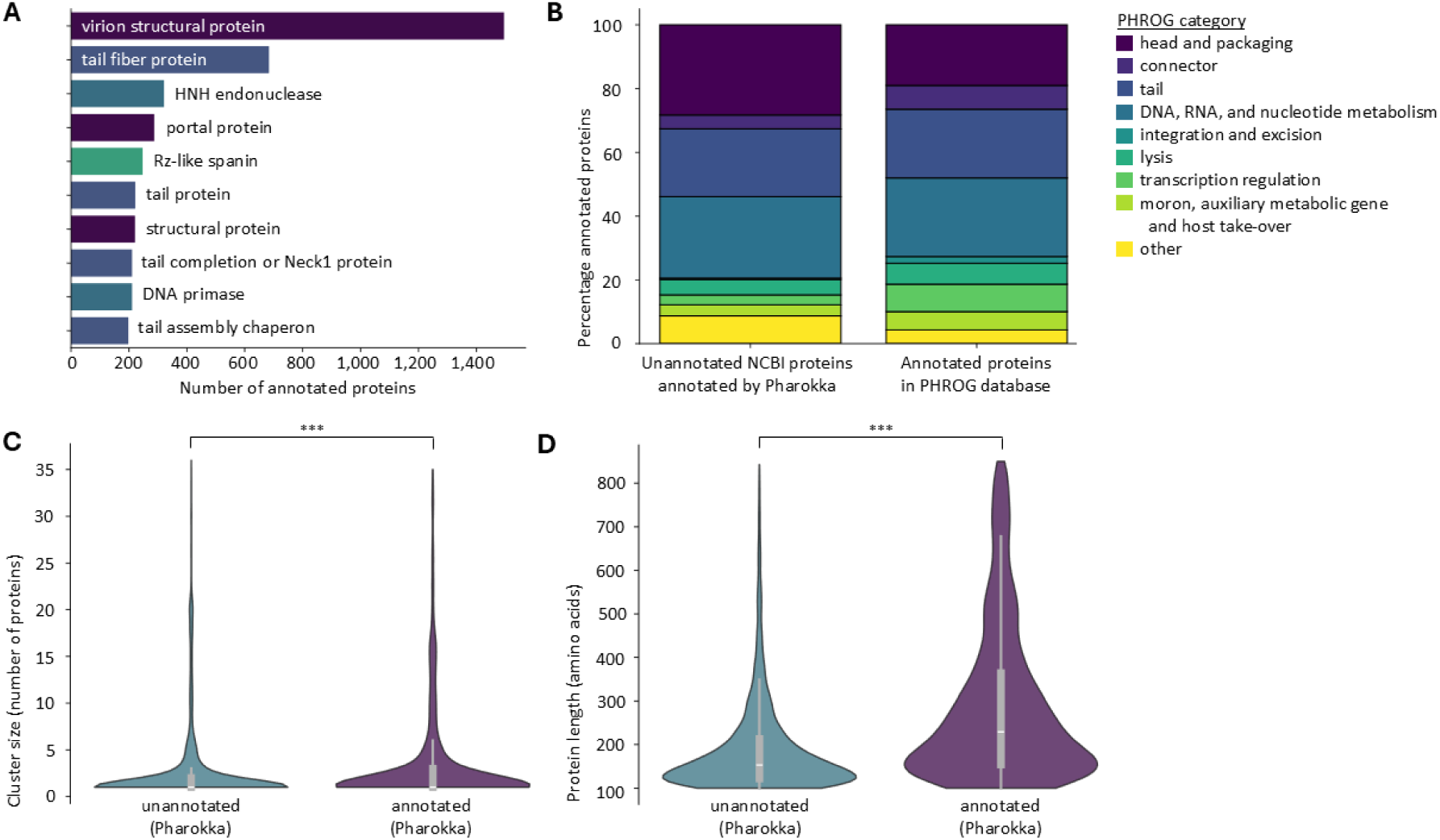
Pharokka annotation of proteins labeled ‘hypothetical/phage protein’ in NCBI protein^29^. **A. Ten most common protein annotations** found by Pharokka^31^. Annotations colored according to PHROG category. **B. Distribution of PHROG categories in the Pharokka annotated proteins, compared to the annotated proteins in PHROG db**^22^**. C. Cluster size of unannotated and annotated protein clusters**, following Pharokka annotation. Violin plot excludes the 1% most extreme values on both ends. *** : p-value of 3.4 x 10^-21^ (one-sided Mann-Whitney U test). **D. Protein length of unannotated and annotated protein clusters**, following Pharokka annotation. *** : p-value of 1.4 x 10^-212^ (one-sided Mann-Whitney U test).

### *Pseudomonas* phage protein structures can be modelled reliably, in spite of their small MSAs and protein sizes

An important prerequisite for structure-informed protein annotation is the availability of high quality protein structures, which were predicted through ColabFold^23^. The majority of the cluster representative proteins was predicted reliably, with mean per-protein pLDDT scores averaging at 74.08 ± 18.10 (Fig. 3A). Strikingly, these results were obtained in the absence of deep MSAs: the median MSA depth was 99 sequences (maximum per-protein MSA depth, Fig. 3B). Indeed, an MSA depth of 10 sequences appeared to be the tipping point for high quality structure predictions (mean per-protein pLDDT score of 82 ± 12 for MSA depths ≥ 10 vs. 55 ± 16 for MSA depths < 10, p = 0.0). Mean per-protein pLDDT increased rapidly with increasing MSA depth when approaching this threshold, but the rate of improvement diminished beyond this point (Spearman’s R^2^ = 0.46, Fig. 3C). Prediction quality did not appear to be correlated to protein length (Pearson’s R^2^ = 0.01, Fig. 3D), although size filtering might have biased those results. Of note, although the majority of cluster representative proteins were smaller than 200 amino acids in length, this did not appear to influence the apparent absence of correlation. Neither binning nor under-/oversampling influenced the observed magnitude of correlation (bin width of 50 amino acids, maximum observed Pearson’s R^2^ = 0.06, average observed Pearson’s R^2^ = 0.02).

**Fig. 3.**
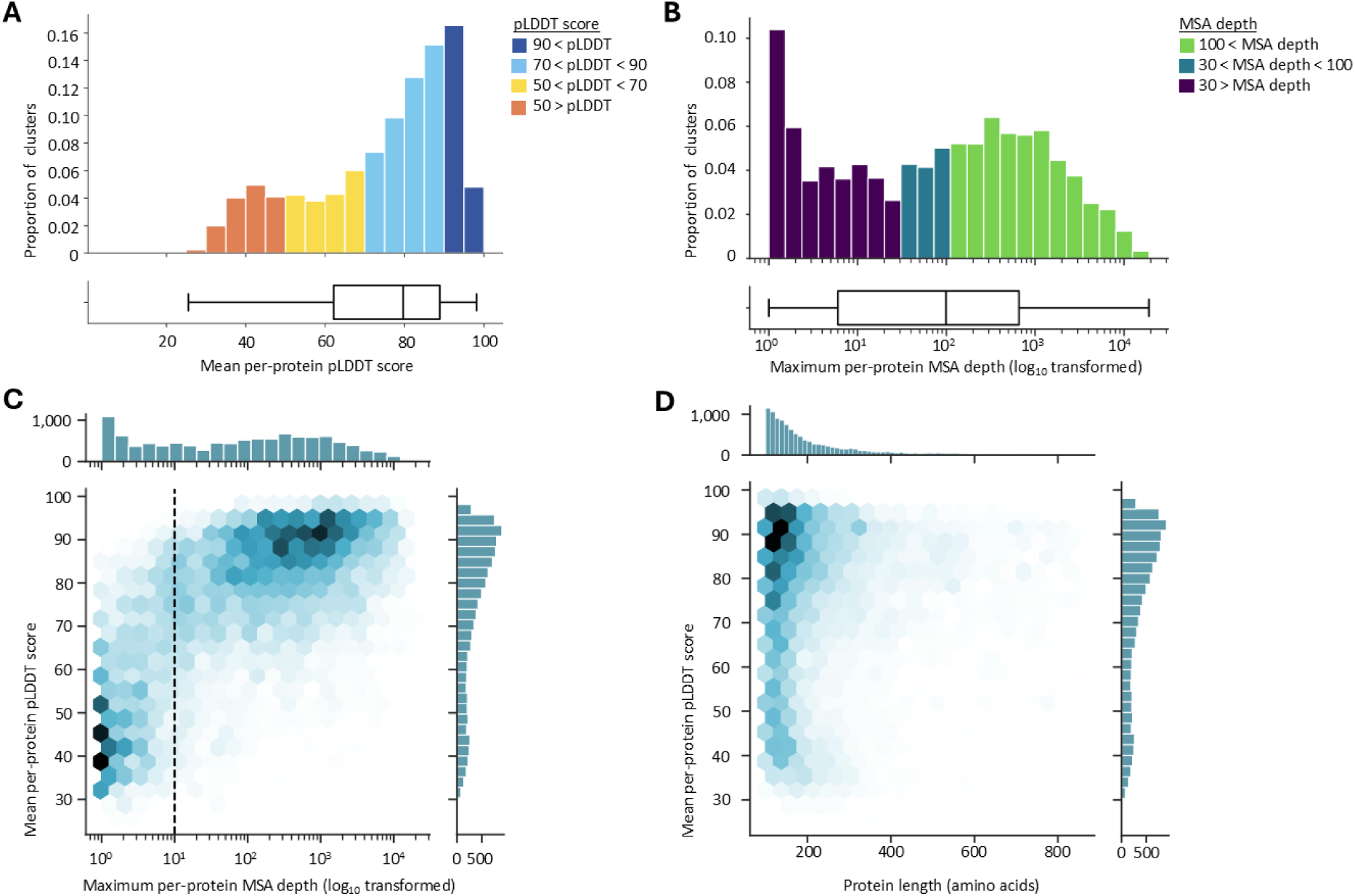
Structure prediction of unannotated phage protein representatives. **A. Mean per-protein pLDDT distribution** of the representative proteins. Bars colored according to pLDDT score. **B. Maximum per-protein MSA depth distribution** of the representative proteins. Bars colored according to MSA depth. **C. Joint distribution of the mean per-protein pLDDT and the maximum per-protein MSA depth** of the representative proteins. Dotted line indicates an MSA depth of 10 sequences. **D. Joint distribution of the mean per-protein pLDDT and protein length** of the representative proteins.

### Phold and AlphaFold databases reveal complementarity in structural similarity to unannotated *Pseudomonas* phage proteins

A second necessary prerequisite for annotation on the basis of structural similarity, is the presence of similar structures in the search database(s). This criterion was met for 69 % of the truly unannotated clusters (representing 87 % of truly unannotated proteins). The phage-specific databases searched by Phold, and the AlphaFold (UniProt50-minimal version) database yielded a relatively high percentage of truly unannotated clusters with at least one significant match (66 and 47 %, respectively, significance cut-off at an E-value of 0.001). The PDB, on the other hand, only significantly matched 9 % of clusters. Although there is substantial overlap between the clusters matching the different databases, the Phold and AlphaFold database appear somewhat complementary (Fig. 4A). While the effect is small, clusters matching none of the three databases represents more genetically diverse proteins, predicted less accurately by ColabFold (Fig. 4B to 4D).

**Fig. 4.**
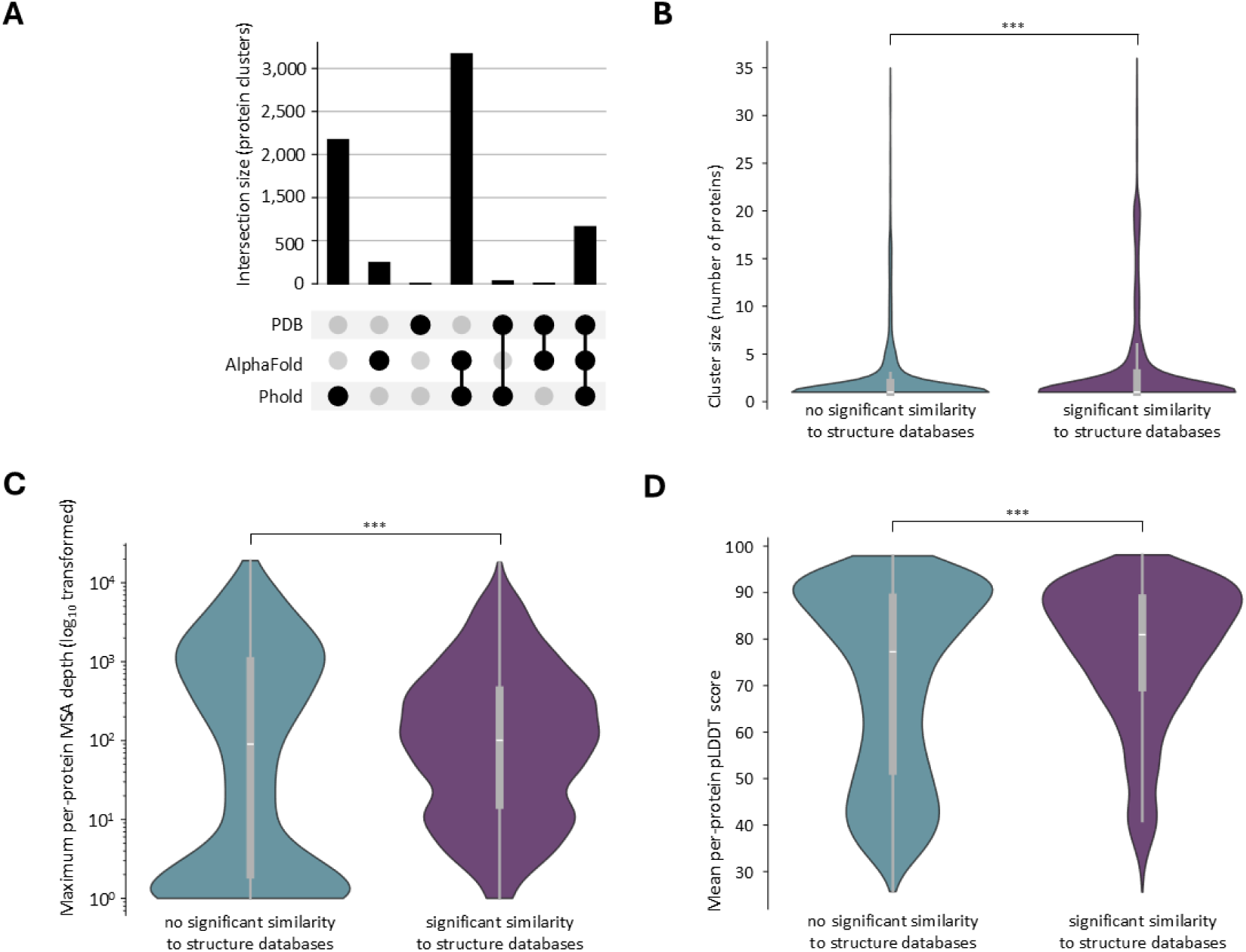
Structural similarity of tuly unannotated phage protein representatives against the PDB, AlphaFold and Phold databases. **A. UpSet plot detailing overlap of structural similarity to the PDB, AlphaFold and/or Phold databases** of the truly unannotated phage protein representatives. Small intersection sizes indicate a lesser degree of complementarity amongst the structure databases. **B. Cluster size of protein clusters with and without significant structural similarity to the PDB, AlphaFold and Phold databases.** Violin plot excludes the 1% most extreme values on both ends. *** : p-value of 1.6 x 10^-21^ (one-sided Mann-Whitney U test). **C. Maximum per-protein MSA depth of protein clusters with and without significant structural similarity to the PDB, AlphaFold and Phold databases.** *** : p-value of 2.8 x 10^-12^ (one-sided Mann-Whitney U test). **D. Mean per-protein pLDDT score of protein clusters with and without significant structural similarity to the PDB, AlphaFold and Phold databases.** *** : p-value of 8.6 x 10^-37^ (one-sided Mann-Whitney U test).

To facilitate downstream annotation, an additional filtering step was implemented, removing poor alignments (alignment lDDT ≤ 0.5, average pLDDT of aligned fraction query ≤ 70) and proteins unlikely to be sharing the same function (probability of belonging to the same SCOP class ≤ 0.5) from the FoldSeek output. This filtering hardly impacted the number of proteins which could be annotated (87.0 % instead of 87.2 % of truly unannotated proteins), but significantly decreased the number of hits to consider during annotation (950,210 vs. 1,483,753 PDB hits, 1,283,164 vs. 1,788,261 AlphaFold database hits).

### ‘Best annotated hit’ method combining PDB, AlphaFold and Phold databases with UniProt annotates 40 % of truly unannotated protein clusters

The first annotation strategy implemented was a ‘best hit’ method, using protein annotations stored in the PDB/AlphaFold database directly. This method resulted in overall low annotation rates, ranging from 6 to 7 % of all truly unannotated protein clusters, and 33 to 46 % of those annotated by Pharokka, for the AlphaFold database and PDB respectively (Fig. 5E and 5F).

**Fig. 5.**
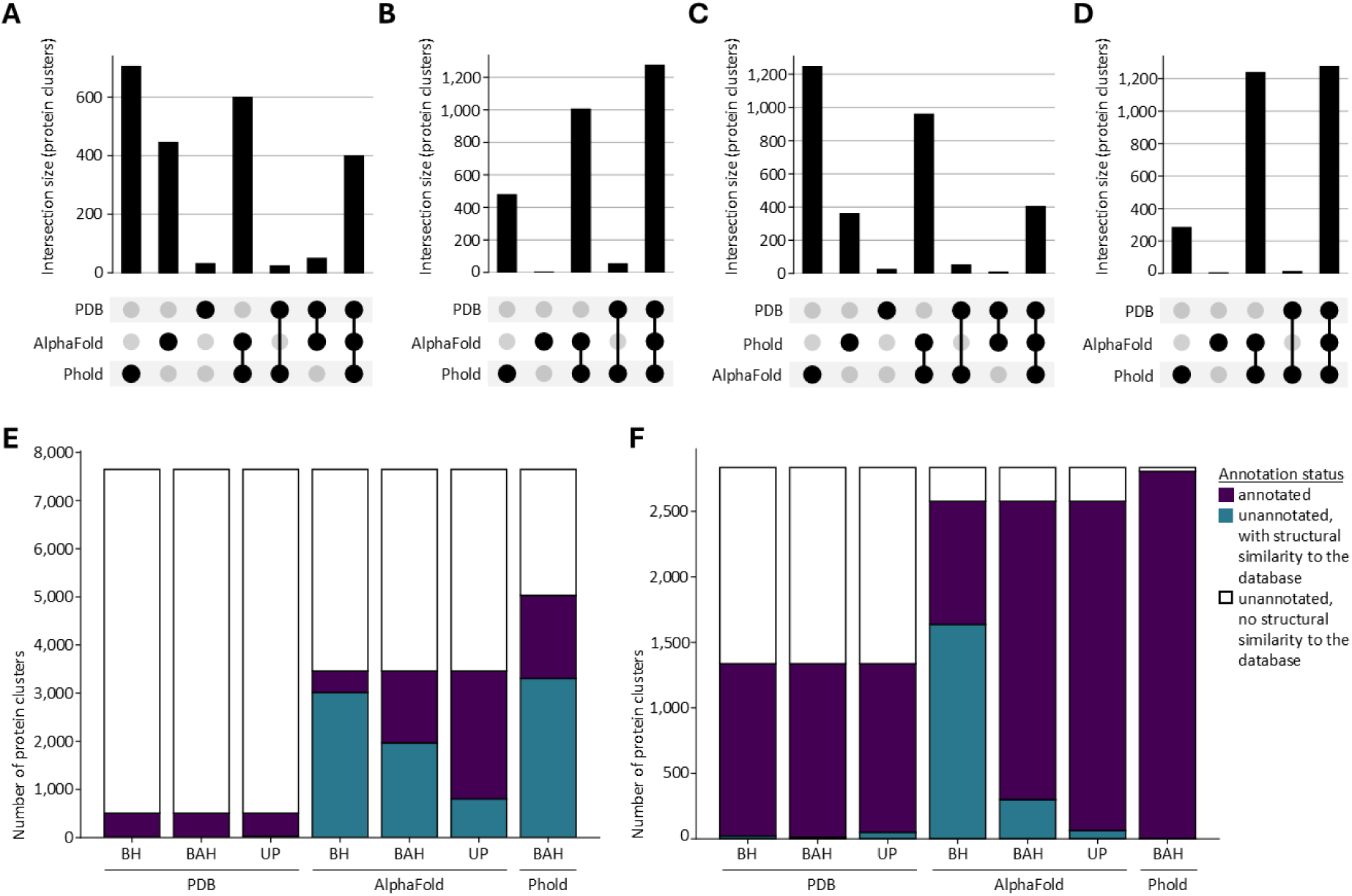
Annotation results of protein clusters across various single hit annotation methods. **A-D. UpSet plots detailing the complementarity in annotated protein clusters** of the subset of truly unannotated protein clusters (A, C) and the subset of those annotated by Pharokka (B, D) using a ‘best annotated hit’ method (A, B), supplemented with UniProt annotations (C, D). **E. Annotation counts of protein clusters considered truly unannotated**, across different methods and databases. Abbreviations: BH: ‘best hit’ – BAH: ‘best annotated hit’ – UP : ‘best annotated hit with UniProt’. Bars colored according to annotation status. **F. Annotation counts of protein clusters previously annotated by Pharokka**, across different methods and databases. Abbreviations: BH: ‘best hit’ – BAH: ‘best annotated hit’ – UP : ‘best annotated hit with UniProt’. Bars colored according to annotation status.

However, as these structure databases contain a considerable number of unannotated proteins, this strategy is unlikely to reach the highest achievable annotation rate. Indeed, a ‘best annotated hit’ strategy, such as implemented in Phold, reached notably higher annotation rates (Fig. 5E and 5F). For the AlphaFold database, annotation rates increased to 20 % of truly unannotated clusters (43 % of those with hits). For the PDB, the annotation rate remained unaffected, as nearly all clusters with hits were already annotated. Phold (in structure comparison mode) reached an annotation rate of 23 % for truly unannotated clusters, and 99 % for those annotated by Pharokka. Jointly, the PDB, AlphaFold and Phold databases annotated 29 % of truly unannotated protein clusters, and 99 % of those annotated by Pharokka (Fig. 5A and 5B). When using UniProt protein names instead of annotations stored in database headers, annotation rates of truly unannotated protein clusters improved even further, to a joint annotation rate of 40 % (fig. 5C).

As expected, annotated proteins were significantly larger, and had better predicted protein structures, built on deeper MSAs (p = 1.0 x 10^-12^, p = 5.3 x 10^-176^ and p = 7.0 x 10^-192^ respectively).

### Three-tiered annotation classification scheme improves quality of annotations

Inspection of the most common annotations found by the ‘best annotated hit’ method, revealed that the most common annotations amongst truly annotated clusters are structural proteins (Fig. 6A and 6B). Moreover, this inspection also clearly illustrated that not all annotations are equally informative, as exemplified by the many “domain of unknown function (DUF)” protein annotations obtained when searching the AlphaFold database. Using an additional tier of annotation classification, we identified 21 % of the ‘best annotated hit’ annotations against the AlphaFold database as being of ‘low information’ (Fig. 6D). Adapting the function assignment module to favor ‘high information’ annotations over ‘low information’ annotations, reduces the percentage of ‘low information’ annotations to 8 % (Fig. 6C and 6D). A similar improvement was observed for the ‘best annotated hit’ method using UniProt annotations, reducing the amount of low informative annotations from 21 to 7 % for the AlphaFold database, while low information annotation rates remained unchanged at 4 % for the PDB .

**Fig. 6.**
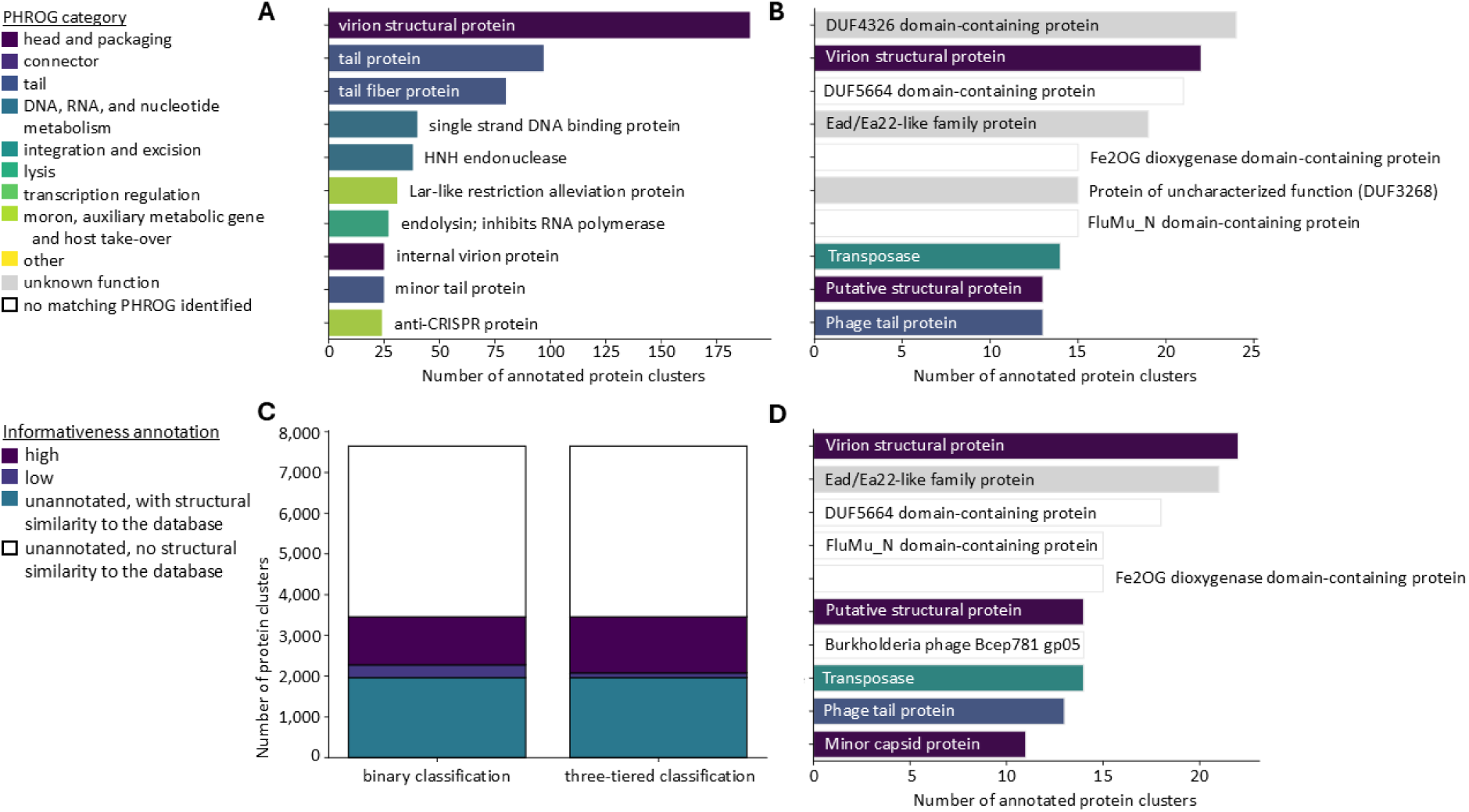
Most common annotations of the truly unannotated clusters, obtained through the best annotated hit method. **A. Ten most common protein cluster annotations found by Phold**. Annotations colored according to PHROG category. **B. Ten most common protein cluster annotations found by the ‘best annotated hit’ method against the AlphaFold database**. Annotations colored according to PHROG category. **C. Ratio of high to low to non-informative protein annotations** among the truly unannotated protein clusters, comparing the binary classification (annotated versus not annotated) and three-tiered classification (high informative versus low informative versus not annotated) scheme. Colored according to annotation informativeness. **D. Ten most common protein cluster annotations found by the ‘best annotated hit’ method against the AlphaFold database using the three-tiered classification scheme**. Annotations colored according to PHROG category.

## Discussion

In this work, we shed light on the viral dark matter of *Pseudomonas* infecting phages, creating a resource of readily available predicted protein structures and annotations. Simultaneously, we investigate the impact of design choices during FoldSeek-based phage annotation efforts, highlighting complementarity across different structure databases and protein annotation resources.

In line with previous findings^6,7,34^, we identified 71% of all proteins from *Pseudomonas* infecting phages in NCBI protein^29^ as unannotated. Of note, NCBI protein is somewhat redundant, possibly inflating the ratio of unannotated proteins. NCBI protein was nevertheless selected as data source, as it provides a more complete picture of the phage protein landscape than UniProt^35^, while remaining a preferred resource among many researchers in the field. A representative, computationally tractable subset of these proteins was extracted, and immediately corrected for the likely overestimation of ‘unannotated’ proteins with Pharokka^31^. Pharokka annotated 27 % of all the ‘unannotated’ protein clusters, which hints at a potentially bigger issue than simply database redundancy. For example, it is possible that protein functions have been elucidated since their initial database deposit, but that these records have not (yet) been updated. Moreover, the incredible genetic diversity and high evolution rate of viral proteins requires detection of remote homology, which due to algorithmic improvements becomes increasingly feasible. Indeed, reannotation with recently updated search databases and progressively powerful annotation tools commonly improves annotation rates^36,37^.

Structure prediction with a ColabFold-like setup revealed accurate structure prediction for the majority of proteins. This is striking, given the shallow MSAs generated and used during structure prediction. These shallow MSAs lie in line with those reported for other viral proteins^14,38^. Structure predictions could potentially be further improved using either additional databases during MSA generation, such as reported by Nomburg and Kim, or the more computationally intensive modelling approach reported by Odai^38,14,15^. While quality of structure prediction proved to be a significant discriminator of ‘annotatable’ proteins, it is yet to be determined whether the reported quality improvements would significantly enhance annotation.

In our subsequent analyses of different variations on FoldSeek-based annotation strategies, we revealed that 31 to 43 % of truly unannotated proteins (29 to 40 % of truly unannotated clusters) could be annotated, when combining hits against PDB, AlphaFold and Phold database. Our results show clear complementarity between these different resources. Although the benefit of using both Phold and AlphaFold databases is most evident, the PDB contains experimental evidence, and remains a preferential structure resource. Using UniProt obtained annotations (through mapping of PDB/AlphaFold database identifiers to UniProt identifiers) rather unexpectedly further improved annotation rates. It must be noted that for many of these annotations, UniProt entries had been removed, and UniParc was consulted to annotate the protein. One could argue that an annotated identical sequence is a good source for annotation, but without investigating the reason for the removal of the initial UniProt entry, caution should prevail.

Although the use of different databases is clearly beneficial, combining and comparing annotations from these resources remains resource intensive and challenging, as neither the PDB, nor the AlphaFold database (or UniProt) use a standardized vocabulary for protein function. To the opposite, PHROGS, which are used by Pharokka and Phold, are highly curated annotations, which leads to more consistent annotations. This discrepancy is clearly observed when looking at the frequency of the most common annotations from the different databases: the most common Phold annotation is shared across over 175 protein clusters, whereas those from the PDB and AlphaFold database are shared by only ∼20 protein clusters. Interestingly, despite the evident complementarity of the different structure database, the most common type of protein annotations is shared: structural proteins. This category was also enriched in the Pharokka annotations. Structural proteins are notoriously divergent and consequently hard to annotate^39–41^, yet an indispensable part of phage biology, which might explain this finding (in addition to the technical bias induced by size filtering).

As expected, the protein clusters which remained unannotated consisted of smaller proteins, which had lower quality structure predictions built on more shallow MSAs. Although this could point to these proteins being more divergent, it is also possible that the lack of detectable homology leads to poor structure prediction, complicating structure similarity based annotation^14^. Furthermore, we cannot exclude the possible presence of spurious proteins in our dataset^42^.

Looking ahead, this resource will be expanded through the removal of size filters, presenting a more complete picture of the viral dark matter, where short proteins, which are generally harder to annotate^43^, are immensely prevalent^44^. Moreover, future analyses will include updating and cross-linking the different annotations across the different resources, to increase our understanding of the most commonly identified annotations. Simultaneously, we will also examine whether ‘single hit’ annotation methods capture the consensus amongst all identified hits, through the use of the pre-existing AlphaFold database structure clusters^45^.

Finally, one should always remain careful when transferring annotations on the basis of similarity^46,47^. As such, it is important to see the annotations provided here as a starting hypothesis, pending further experimental validation and characterization. Nonetheless, we foresee this resource as a potent catalysator of future research and applications, ranging from facilitating phage therapy (through more completely predicted phage capabilities), to providing SynBio with more genetically diverse building blocks (such as depolymerases that remained previously unidentified)^34^.

## Materials and methods

### Included phages and proteins

An overview of the different *Pseudomonas* infecting phage species was obtained by downloading all entries of the Virus-Host database (version February 25, 2024^32^) that list ‘*Pseudomonas*’ in the ‘host’ field. Protein information was fetched from NCBI Protein^29^ through eutils (accessed February 26^th^, 2024). Phages for which NCBI protein did not contain any entries were removed from the dataset. Unannotated proteins were obtained by filtering the NCBI protein names on ‘hypothetical protein’ and/or ‘phage protein’. Only those with a size between 100 and 850 amino acids were retained.

A representative set of phage proteins of unknown function was generated using MMSeqs2^33^ v.14.7e284. Briefly, all obtained phage proteins of unknown function were clustered at 90% sequence identity and 80% bidirectional coverage using MMseqs2 ‘cluster’. A representative protein for each cluster was then obtained using MMseqs2 ‘createsubdb’.

### Structure prediction and structural similarity search

Protein structures were predicted with VIBFold^48^ (non-default settings: --do-relax best), an adapted version of AlphaFold2^10^ v2.3.1 that utilizes MSAs generated through the MMseqs2 API^33^, inspired by ColabFold^23^. Next, predicted protein structures were queried for structural similarity with FoldSeek^19^ v.8.ef4e960 using ‘easy-search’ against databases ‘PDB’ v.2024-02-20 and ‘Alphafold/UniProt50-minimal’ v.4 (non-default settings: --format-output "query, target, fident, alnlen, mismatch, gapopen, qstart, qend, tstart, tend, evalue, bits, prob, lddt, lddtfull"). FoldSeek hits were then filtered on an E-value < 0.001.

### Protein annotation with Pharokka

The representative set of phage proteins of unknown function was annotated with Pharokka v.1.7.5^31^ (databases v.1.4.0^49^) using ‘pharokka_proteins.py’. Specifically, functional annotations were generated by matching each protein sequence to the PHROGs^22^, VFDB^20^ and CARD^24^ databases using MMseqs2^33^ and PyHMMER^50^.

### Protein annotation with Phold

The representative set of phage proteins of unknown function was annotated with Phold v.0.2.0^17^ (databases v.0.2.0^51^) using ‘proteins-compare’. Specifically, functional annotations were generated by matching each protein structure to the PHROGs^22^, VFDB^20^ and CARD^24^, DefenseFinder^27^, acrDB^21,26^, Netflax^25^ databases using FoldSeek^19^.

### Protein annotation with FoldSeek using the PDB and AlphaFold database

FoldSeek hits were further filtered on alignment lDDT > 0.5, average pLDDT of aligned portion of query > 70, and prob(ability of belonging to the same SCOP class) > 0.5, then sorted by bitscore.

In the ‘best hit’ method, functional annotations were generated by matching the first hit to its annotation stored in the PDB/AlphaFold database header file. Annotated hits were discriminated from non-annotated hits using a set of self-defined regular expressions.

In the ‘best annotated hit’ method, functional annotations were generated by matching the first annotated hit (according to the previously defined regular expressions) to its annotation stored in the PDB/AlphaFold database header file.

In the ‘best annotated hit’ method supplemented with information from UniProt, protein names were obtained using the PDB data API^52^ (accessed July 15th, 2024) and UniProt API^35^ (UniProt release 2024_03).

For the ‘best annotated hit’ method using a three-tiered annotation classification, additional regular expressions were defined to discriminate low informative annotations from proper annotations. During annotation, proper annotations were favored over low informative annotations, which were in turn favored over the absence of a functional annotation.

### Data visualization

Sankey plots visualizing phage and bacterial taxonomy were created with Pavian^53^, upset plots with UpSetPlot v.0.9.0^54^. All other plots were generated with Seaborn v.0.13.2^55^.

### Statistical analysis

Throughout this work, statistical analyses were implemented through SciPy v.1.16.0^56^, using ‘pearsonr’ and ‘spearmanr’ for correlation analysis, and ‘mannwhitneyu’ for comparison of distributions. Over- and undersampling was performed with ‘Imbalanced-learn’ v.0.13.0^57^.

## Data availability

Predicted protein structures, FoldSeek output files, functional annotations and the cluster assignment table generated in this study have been deposited in Zenodo under doi: 10.5281/zenodo.16033188.

## Code availability

All code related to this study is available under an MIT license through GitHub at https://github.com/hannelorelongin/FoldFirstAskLater.

## Acknowledgements

We thank dr. Jasper Zuallaert (Center for Medical Biotechnology, VIB, Belgium) for his implementation of and help with VIBFold.

This research was financially supported by the KU Leuven C1 project ACES [C16/20/001] (H.L., H.H., R.L., V.v.N.), by the Research Foundation - Flanders (FWO, Belgium) ‘long stay abroad’ grant [V412225N] (H.L.), by the National Institutes of Health (NIH) National Institute of Diabetes and Digestive and Kidney Diseases [RC2DK116713] (R.A.E), and the Australian Research Council [DP220102915; DP250103825] (R.A.E.). H.L. holds a PhD fellowship fundamental research [11PCC24N] of the Research Foundation - Flanders (FWO, Belgium).

The computational resources and services used in this work were provided by the VSC (Flemish Supercomputer Center), funded by the Research Foundation - Flanders (FWO) and the Flemish Government.

## Declarations

The authors declare no competing interests.

